# The pivotal role of a novel free fatty acid receptor GPR164 in the intestinal barrier function

**DOI:** 10.1101/2025.02.20.639380

**Authors:** Takako Ikeda, Yuki Masujima, Keita Watanabe, Akari Nishida, Mayu Yamano, Miki Igarashi, Nobuo Sasaki, Hironori Katoh, Ikuo Kimura

**Affiliations:** Laboratory of Molecular Neurobiology, Graduate School of Biostudies, Kyoto University, Sakyo-ku, Kyoto 606-8501, Japan; Department of Molecular Endocrinology, Graduate School of Pharmaceutical Sciences, Kyoto University, Sakyo-ku, Kyoto 606-8501, Japan; Southern TOHOKU Research Institute for Neuroscience, 255 Furusawa, Asao-ku, Kawasaki, Kanagawa 215-0026, Japan; Laboratory of Mucosal Ecosystem Design, Institute for Molecular and Cellular Regulation, Gunma University, Maebashi, Japan; Department of Biological Chemistry, Graduate School of Science, Osaka Metropolitan University, Gakuen-cho, Naka-ku, Sakai, Osaka 599-8531, Japan

## Abstract

GPR164 is a novel free fatty acid receptor, activated by both short-chain fatty acids and medium-chain fatty acids, and expressed throughout the gastrointestinal tract. Although GPR164 is reported to be involved in the release of gut hormones, the physiological functions of this receptor in the maintenance of intestinal homeostasis remain unclear. In this study, we explored the role of GPR164 in regulating intestinal barrier function using mice lacking *Gpr164* gene (*Gpr164*^-/-^). A loss-of-function mutation in *GPR164* promoted cell proliferation and disrupted the intestinal barrier function in both Caco-2 cell line and mice. Genome-wide RNA-seq analysis revealed that *GPR164* deletion caused aberrant wnt/β-catenin signaling, and the intraperitoneal injection of wnt/β-catenin inhibitor ameliorated a series of abnormalities of *Gpr164*^-/-^ mice. *Gpr164*^-/-^ mice also exhibited gut microbial dysbiosis and severe inflammation, indicating that deletion of *Gpr164* causes similar pathologies observed in patients with inflammation bowel disease (IBD). Thus, our findings uncover the pivotal role of GPR164 in the maintenance of intestinal barrier function, providing an attractive clinical target for IBD.

## Introduction

GPR164, also known as Olfr558 in mouse and OR51E1 in human, was initially identified as an olfactory chemosensory receptor. Mouse Olfr558 and human OR51E1 are encoded by *Or51e1* and *OR51E1* gene, respectively, and expressed in several tissues including the olfactory epithelium. In peripheral tissues, GPR164 is activated by free fatty acids (FFAs) such as butyrate and nonanoate, thereby being considered as a free fatty acid receptor (FFAR) ^1,2^. Although a large number of FFAs such as iso-butyrate, valerate and nonanoate has been identified as ligands for GPR164, physiological abundance of these FFAs is not enough to activate GPR164 in the intestine. Contrary to these FFAs, butyrate is usually present in the colon at millimolar level, and activates GPR164 with an EC_50_ value of 0.51 mM^3^. Therefore, it is plausible that GPR164 is preferentially activated by butyrate in colon and plays an important role in regulating the intestinal homeostasis. Recently, GPR164 has been reported to have a potential relevance for the entero-endocrine signaling. In pigs, OR51E1 is expressed throughout the gastrointestinal tract, and co-localized with peptide YY (PYY)- and 5-hydroxytriptamine (5-HT)-expressing cells^4^. Furthermore, in human enteroendocrine L-cell line NCI-H716, OR51E1 is co-localized with glucagon-like peptide-1 (GLP-1), and nonanoate increases the secretion of GLP-1 and PYY through activation of this receptor^5^. These results suggest an important role for GPR164 in the regulation of gut hormone release. However, the physiological functions of GPR164 in the maintenance of intestinal homeostasis remain largely unclear.

Butyrate is a free fatty acid classified into the short-chain fatty acid (SCFA), and plays an important role in many physiological functions such as energy metabolisms and immune responses^6,7^. SCFAs are bioactive end-products produced by the gut microbiota through anaerobic fermentation in the colon^8^. Altered microbial composition (dysbiosis) affects the production of SCFAs, thereby contributing to various diseases such as obesity, type 2 diabetes mellitus (T2DM), and inflammatory bowel disease (IBD). SCFAs are thus crucial mediator linking gut microbiota and optimal host health. In the intestinal epithelial cells, butyrate serves as an immunomodulator with anti-inflammatory effects. Butyrate suppresses pro-inflammatory cytokine production (e.g., tumor necrosis factor-alpha) in part through inhibiting NF-κB activation, and ameliorates the development of colitis by promoting differentiation of regulatory T cells^9,10^. Administration of butyrate in patients with ulcerative colitis alleviates intestinal inflammation, suggesting a potential therapeutic agent for the treatment of IBD^11^. In addition, butyrate enhances the intestinal barrier function by up-regulating the expression of tight junction and mucin proteins, conferring increased protection against enteric toxins and pathogens^12,13^. However, the underlying mechanisms of butyrate-mediating maintenance of the intestinal barrier function remain elusive.

The intestinal epithelial cells self-renew rapidly, which allows for exerting multiple functions involving digestive and absorptive activities^14^. This characteristically rapid turnover relies on the highly proliferative stem cells. Intestinal stem cells in the crypt give rise to transit amplifying cells, which gradually differentiate into other epithelial lineages such as absorptive enterocytes and secretory cells (e.g. enteroendocrine cells). These differentiated cells bidirectionally migrate along the crypt-villus axis, and are removed by apoptosis upon reaching the villus tip. Wnt/β-catenin signaling pathway (Wnt signaling) regulates cell fate along the crypt-villus axis. Activation of Wnt signaling in the crypt promotes cell proliferation, while attenuation of this signaling in the villus results in cell cycle arrest. Aberrant activation of Wnt signaling initiates and develops colorectal cancers, suggesting an essential role of Wnt signaling in maintaining intestinal homeostasis. Intriguingly, GPR164 is overexpressed in human primary prostate cancers and prostate cancer cell line LNCaP, and used as a biomarker of human prostate cancer^15^. Treatment with nonanoate, GPR164 agonist, suppresses the proliferation of LNCaP cells and induces cellular senescence^16^. This cytostatic effect induced by activation of GPR164 is mediated by p38 MAPK signaling pathway, implying an essential role of GPR164 in the cell cycle regulation. As well as in the prostate cancer, high expression of GPR164 is observed in lung carcinoids and digestive neuroendocrine carcinomas^17,18^. Therefore, GPR164 has gained as a therapeutic target for the treatment of cancers.

In this study, we investigated the physiological importance of GPR164 as a key factor for regulating intestinal barrier function using *Gpr164*^-/-^ mice. The endogenous functions of this receptor remain obscure, and the underlying mechanisms for the maintenance of intestinal homeostasis are not fully understood. Therefore, our findings provide important insight into developing therapeutic targets for the treatment of IBD and cancers.

## Results

### Loss of *OR51E1* results in increased cell cycle progression and intestinal barrier dysfunction in Caco-2 cells

In peripheral tissues, GPR164 stimulates adenylyl cyclase in response to FFAs and increases intracellular cAMP levels. Although many FFAs have been identified as ligands of GPR164, some of these FFAs are not suited to act as an endogenous ligand in the intestine. Therefore, we first performed ligand screening assay using Or51e1-overexpressing HEK293 cells. To functionally express transfected Or51e1 on the cell surface, receptor-transporting proteins (RTPs) was co-transfected with Or51e1 because GPR164 is an olfactory receptor that requires the chaperones such as RTPs for the ectopic expression in other non-nasal tissues^19^. In mammals, RTP family consists of 4 members (RTP1-4), and a shorter form of RTP1 (RTP1S) was reported to promote cell-surface expression of GPR164^3^. However, mRNA expression level of *Rtp1S*, *Rtp2* and *Rtp3* in mouse colon were lower than that of *Rtp4* (Fig.1a), suggesting that RTP4 is a dominant chaperone required for GPR164 trafficking to the plasma membrane in the colon. The Or51e1 expression was confirmed by transient co-transfection with either RTP1S or RTP4 in HEL293 cells (Supplementary Fig. 1a), and intracellular cAMP levels were up-regulated by the treatment with butyrate or valerate (Fig. 1b). It is interesting to note that decanoate increased cAMP level only when cells were co-expressed with RTP1S (Fig. 1b), indicating that SCFAs, especially butyrate, play central roles in regulating biological processes through GPR164 activation in the colon. We next investigated the functions of GPR164 using *OR51E1* knockout Caco-2 (*OR51E1* KO) cells generated by the CRISPR/Cas9 system (Supplementary Fig. 1b). GPR164 was previously reported to induce cell cycle arrest and cell death^20^. Therefore, we examined the mRNA expression levels of genes associated with cell cycle regulation, and found the decreased expression of *p21* and *p27* in *OR51E1* KO cells at any time points (Fig. 1c). Additionally, the mRNA expression of cyclin genes in *OR51E1* KO cells was lower until reaching confluence (day 0-9) but higher after confluence (day 12-18) than that in WT cells (Fig. 1c). Consistent with these results, *OR51E1* KO cells exhibited increased cell growth after day 15, but not until day 12 when compared to WT cells (Fig. 1d), suggesting that GPR164 is a key regulator of cell proliferation. Caco-2 cells, which are derived from a colon carcinoma, form a polarized epithelial cell monolayer after reaching confluence, and are used as a model of the intestinal epithelial barrier. In *OR51E1* KO cells, mRNA expression of *Occludin* and *Claudin-3* was decreased, whereas *Zo-1* mRNA expression was increased after confluence compared with WT cells (Fig. 1e). In addition, reduction in transepithelial electrical resistance (TER), a reliable in vitro measurement of the barrier function, was observed in *OR51E1* KO cells after day 12 (Fig. 1f). Although GPR164 was reported to induce p53-dependent cell cycle arrest and apoptosis^20^, we could not detect a change in p53 protein expression between WT and *OR51E1* KO cells (Fig. 1g and Supplementary Fig. 1c). Taken together, these results suggest that GPR164-mediated cell cycle regulation plays an important role in maintaining the intestinal barrier functions in Caco-2 cells.

**Fig. 1.**
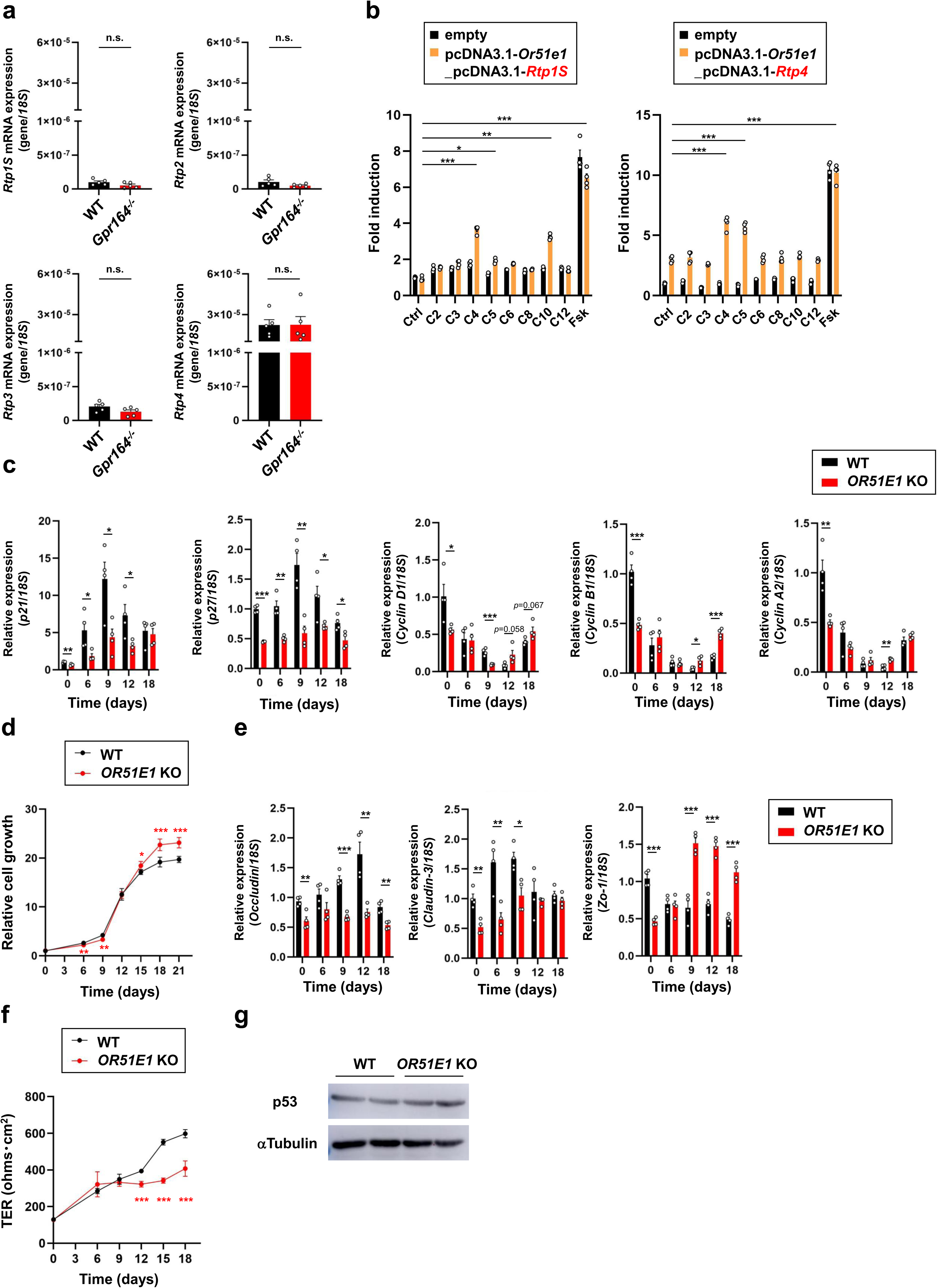
O*R*51E1-knockout Caco-2 cells exhibit the intestinal barrier dysfunction. (a) The mRNA expression levels of RTP family members determined by qRT-PCR. Total RNA was extracted from colon tissue of WT and *Gpr164^-/-^* mice (n=5). Error bars represent the mean ±SEM. n.s., not significant (Student’s *t*-test). (b) Measurement of intracellular cAMP in Or51e1-overexpressing HEK293 cells. Cells were co-transfected with HA-tagged Or51e1 and receptor-transporting protein (left; RTP1S, right; RTP4), and treated with the indicated FFA at 1 mM for 10 mins. Folskolin (FSK) was used as a positive control. The values represent the mean ±SEM from two independent experiments (n=4). \**P* < 0.05, \*\**P* < 0.01, \*\*\**P* < 0.001 (Dunn’s test; left, Dunnett’s test; right). (c) The mRNA expression levels of cell cycle genes determined by qRT-PCR. The bars represent the relative mRNA levels from at least three independent experiments (n=4). Error bars show the mean ±SEM. \**P* < 0.05, \*\**P* < 0.01, \*\*\**P* < 0.001 (Student’s *t*-test). (d) Measurement of cell proliferation by crystal violet staining. Cultured cells were stained with crystal violet at the indicated times, and absorbance was measured at OD_590_ nm. The values represent the mean ±SEM from three independent experiments (n=5). \**P* < 0.05, \*\**P* < 0.01, \*\*\**P* < 0.001 (Student’s *t*-test). (e) The mRNA expression levels of tight junction markers determined by qRT-PCR. The bars represent the relative mRNA levels from at least three independent experiments (n=4). Error bars show the mean ±SEM. \**P* < 0.05, \*\**P* < 0.01, \*\*\**P* < 0.001 (Student’s *t*-test). (f) Monitoring of TER at different times. The values represent the mean ±SEM from at least three independent experiments (n=9). \*\*\**P* < 0.001 (Mann-Whitney *U*-test). (g) Representative image of p53 protein expression. Total cell lysates extracted from WT or *OR51E1* KO cells were subjected to immunoblot analysis using an anti-p53 or anti-αTubulin antibody.

### *Gpr164*^-/-^ mice exhibit colonic hyperplasia and impaired barrier function

Next, we investigated the physiological expression levels of *Or51e1* in the mouse intestine and olfactory epithelium. Consistent with previous reports^1^, *Or51e1* was expressed not only in the olfactory epithelium but also in the intestine (Fig. 2a). Although *Or51e1* expresses throughout the intestine, higher expression was observed in the colon (Fig. 2a). We therefore examined the morphological changes in *Or51e1* gene knockout (*Gpr164^-/-^*) mice by using paraffin-embedded cross-section of colon (Supplementary Fig. 2a, b, c). Hematoxylin and eosin (HE) staining revealed increased area of the colon in *Gpr164^-/-^*mice (Fig. 2b), and whole length of the colon and the length of colonic glands in *Gpr164^-/-^*mice were longer than those in WT mice (Fig. 2c, d). To examine whether the colonic hyperplasia observed in *Gpr164^-/-^* mice was caused by aberrant cell cycle progression, we examined the expression of cell cycle genes and Ki67 cell proliferation marker protein. In *Gpr164^-/-^* mice, the mRNA expression of *p21* and *p27* was decreased, but that of cyclin genes was increased (Fig. 2e). Immunohistochemical analysis of Ki67 confirmed enhanced cell proliferation in *Gpr164^-/-^* mice (Fig. 2f), suggesting that accelerated cell proliferation caused by *Gpr164* deficiency leads to the colonic hyperplasia. Furthermore, decreased expression of tight junction marker genes was observed in *Gpr164^-/-^* mice (Fig. 2g). We thus assessed the role of GPR164 in the intestinal barrier function *in vivo* by using fluorescein isothiocyanate (FITC) labeled-dextran. After oral administration of FITC-dextran, plasma level of FITC-dextran in *Gpr164^-/-^*mice was elevated (Fig. 2h). These findings are consistent with the results obtained *in vitro* model cells, and suggest an essential role of GPR164 for maintaining the intestinal barrier function.

**Fig. 2.**
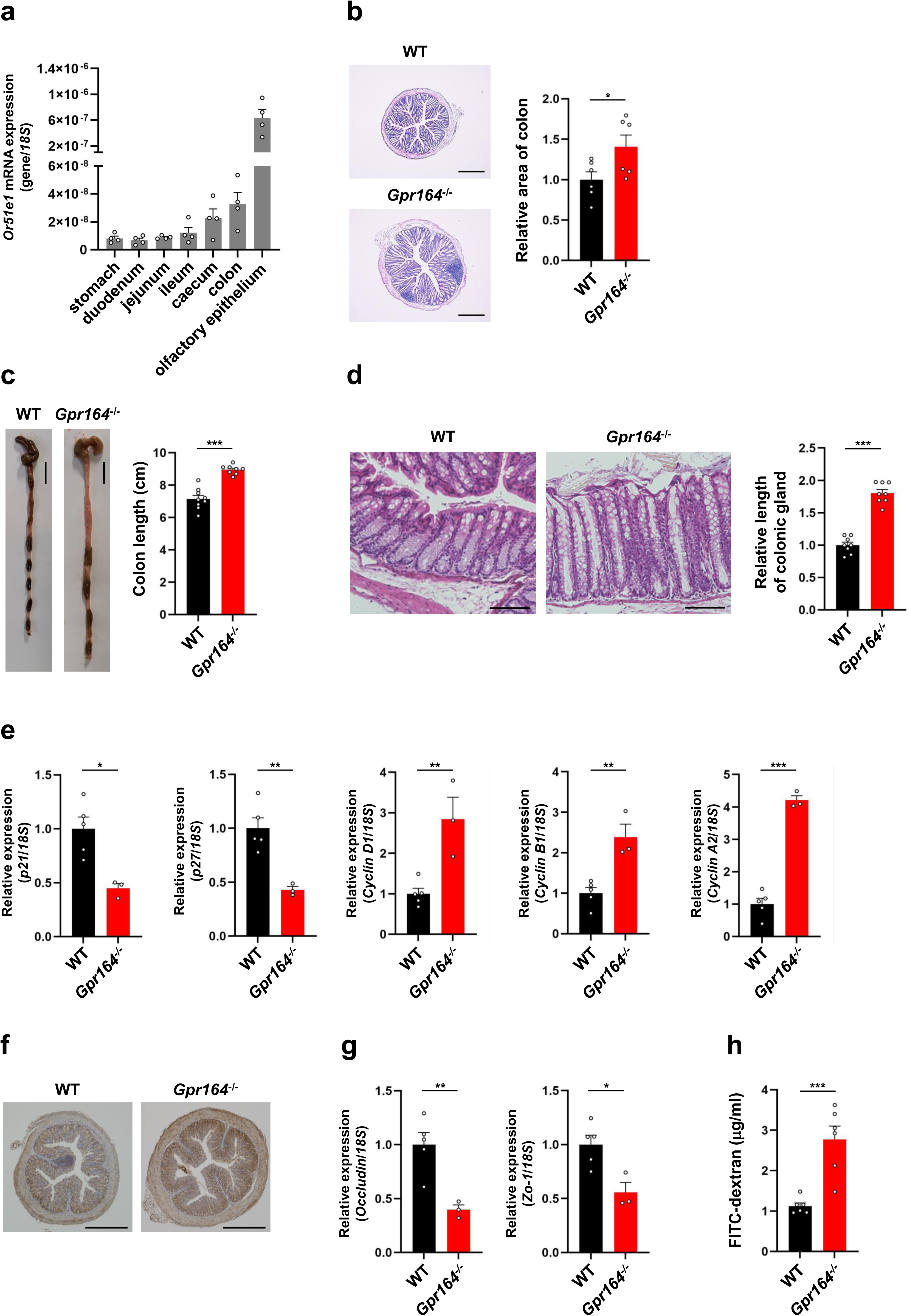
G*p*r164^-/-^ mice exhibit colonic hyperplasia and intestinal barrier dysfunction. (a) *Or51e1* mRNA expression determined by qRT-PCR. Total RNA was extracted from gastrointestinal tract of WT mice (n=4). Error bars represent the mean ±SEM. (b) Representative images of colon sections stained with hematoxylin-eosin (HE). The cross-sections of colon obtained from WT and *Gpr164^-/-^* mice were stained with HE, and the area of colon was measured using ImageJ software (n=6). Scale bar, 500 μm. Error bars represent the mean ±SEM. \**P* < 0.05 (Student’s *t*-test). (c) Representative images of colon (n=8-9). Scale bar, 1 cm. Error bars represent the mean ±SEM. \*\*\**P* < 0.001 (Student’s *t*-test). (d) The colon sections stained with HE. The length of colonic gland was measured using ImageJ software (n=8). Scale bar, 100 μm. Error bars represent the mean ±SEM. \*\*\**P* < 0.001 (Student’s *t*-test). (e) The mRNA expression levels of cell cycle genes determined by qRT-PCR. Total RNA was extracted from colon tissue of WT and *Gpr164^-/-^* mice (n=3-5). Error bars represent the mean ±SEM. \**P* < 0.05, \*\**P* < 0.01, \*\*\**P* < 0.001 (Student’s *t*-test). (f) Representative images of immunohistochemical staining for Ki67 in colon. The cross-sections of colon obtained from WT and *Gpr164^-/-^* mice were stained with anti-Ki67 antibody. DAB was used for the detection of Ki67 protein, and hematoxylin was used for nuclei staining (n=5). Scale bar, 500 μm. (g) The mRNA expression levels of tight junction markers determined by qRT-PCR. Total RNA was extracted from colon tissue of WT and *Gpr164^-/-^* mice (n=3-5). Error bars represent the mean ±SEM. \**P* < 0.05, \*\**P* < 0.01 (Student’s *t*-test). (h) Assessment of intestinal permeability by measuring plasma levels of FITC-dextran. The fluorescence intensity of FITC-dextran in plasma was measured (n=6). Error bars represent the mean ±SEM. \*\*\**P* < 0.001 (Student’s *t*-test).

### Loss of *Gpr164* induces gut microbial dysbiosis and intestinal inflammation

The mucus layer, which mainly composed of Muc2 mucin *O*-glycan, serves as a physical barrier against luminal pathogens in the colon^21^. To investigate whether the impairment of the intestinal barrier function observed in *Gpr164^-/-^* mice is caused by the defects in the colon mucus, alcian blue-periodic acid Schiff (PAS) staining was performed. The colon section from *Gpr164^-/-^* mice was strongly stained with alcian blue-PAS, but the mucus layer of *Gpr164^-/-^* mice was thinner than that of WT mice (Fig. 3a, b). The gut microbiota inhabits the colon mucus which separates bacteria and host (physical barrier) and contains anti-bacterial proteins responsible for blocking the bacteria from penetrating the epithelial cell barrier (immune barrier). Therefore, impaired mucus barrier function alters the microbial community and induces colonic inflammation^22^. Given that GPR164 plays an important role in the mucus barrier function, we examined the alterations in gut microbial composition by using fecal samples of WT or *Gpr164^-/-^* mice. 16S rRNA sequencing showed that the relative abundances of bacterial phyla were markedly changed between WT and *Gpr164^-/-^* mice (Fig. 3c). Notably, the microbial composition of *Gpr164^-/-^* mice shifted toward that of WT mice fed high-fat diet (HFD), a well-known risk factor for metabolic and inflammatory diseases induced by the gut microbial dysbiosis (Fig. 3c). The hierarchical clustering analysis also confirmed marked alterations in the microbial composition with decreased abundance of Bacteroidota and increased abundance of Firmicutes in *Gpr164^-/-^* mice (Fig. 3d, e). The Firmicutes/Bacteroidota (F/B) ratio represents a biomarker of gut microbial dysbiosis associated with inflammation. Therefore, we next examined the involvement of GPR164 in inflammation. Long-term HFD feeding dramatically increased plasma TNFα level in WT mice, and *Gpr164* deficiency also caused severe inflammation (Fig. 3f). Notably, plasma level of TNFα in *Gpr164^-/-^* mice was comparable to that observed in WT mice-fed HFD, and *Gpr164^-/-^* mice exhibited severe inflammation even under normal diet conditions. We further examined the impact of intestinal dysbiosis on the production of microbial metabolites, and found the decreased levels of butyrate in the colon of *Gpr164^-/-^*mice (Fig. 3g). Taken together, these results indicate that loss of *Gpr164* causes the mucus barrier dysfunction, which leads to gut microbial dysbiosis associated with decreased production of butyrate in the colon. The disruption of intestinal barrier function regulated by GPR164 results in severe inflammation, suggesting a pivotal role of GPR164 in maintaining intestinal homeostasis.

**Fig. 3.**
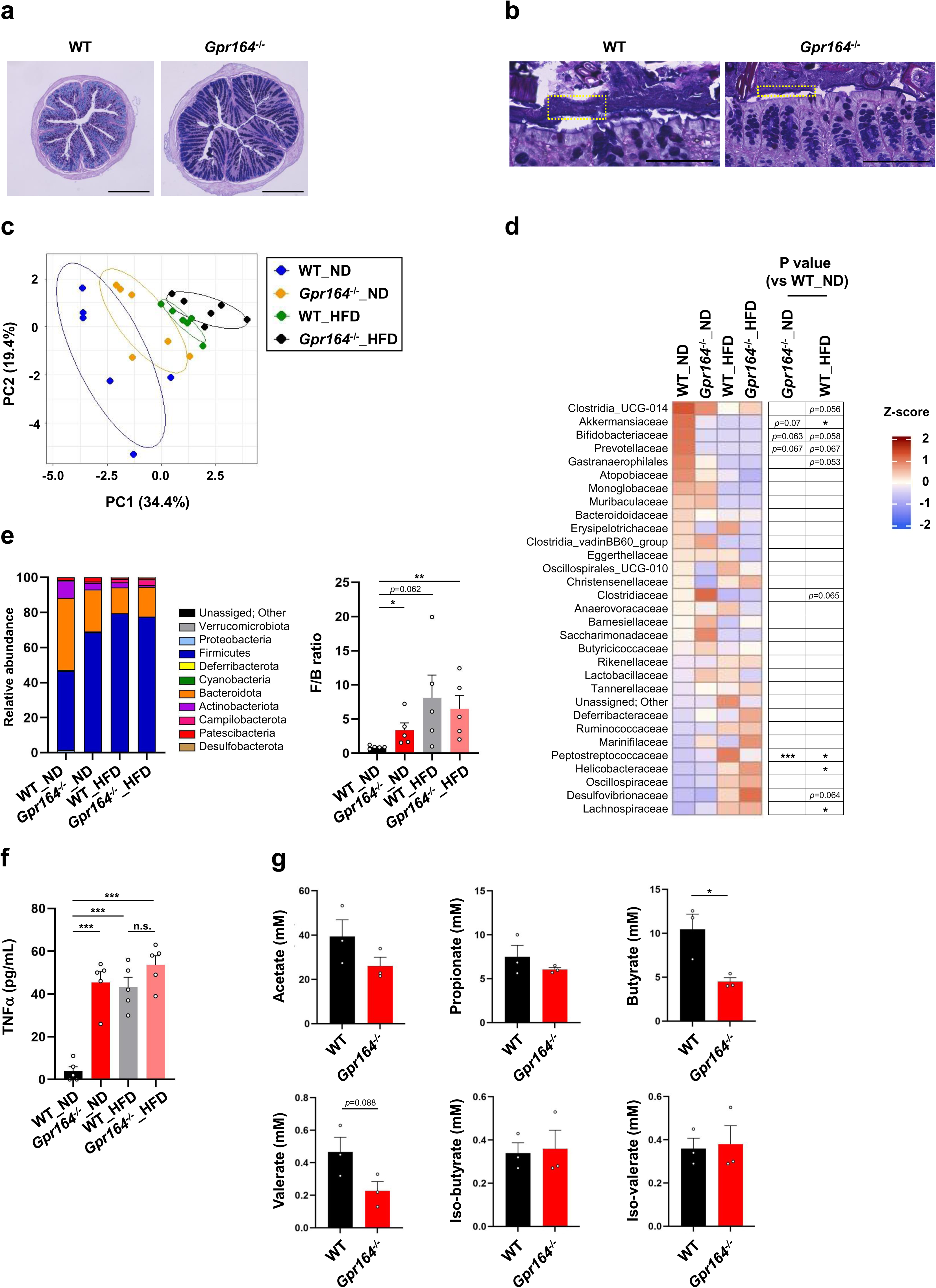
Loss of *Or51e1* results in the gut microbial dysbiosis and inflammation. (a, b) Representative images of colon sections stained with alcian blue-periodic acid Schiff (PAS) (n=5). High- (a) or low- (b) magnification cross-sectional images of colon obtained from WT and *Gpr164^-/-^* mice are shown. A part of mucus layer was bounded by yellow dotted lines (b). Scale bar, 500 μm (a) and 100 μm (b). (c) Principal coordinate analysis of fecal microbiota from WT and *Gpr164^-/-^* mice-fed either a normal diet (ND) or a high-fat diet (HFD) for 10 weeks (n=6). (d) Heatmap of relative abundance of taxonomic units in the gut microbiota at the family level (n=6). (e) Relative abundance of taxonomic units in the gut microbiota at the phylum level (left), and Firmicutes/ Bacteroidota (F/B) ratio (right) (n=5). Error bars represent the mean ±SEM. \**P* < 0.05, \*\**P* < 0.01 (Student’s *t*-test). (f) Plasma levels of TNFα in WT and *Gpr164^-/-^*mice-fed either a ND or a HFD for 10 weeks (n=5). Error bars represent the mean ±SEM. \*\*\**P* < 0.001, n.s., not significant (Tukey-Kramer test). (g) SCFA levels in colonic contents (WT; n=3, KO; n=5). Error bars represent the mean ±SEM. \**P* < 0.05 (Student’s *t*-test).

### *Gpr164*^-/-^ mice has defects in the intestinal epithelial lineage caused by aberrant Wnt signaling

Mucins are condensed in the secretory granules of goblet cells. Considering that the colon sections from *Gpr164^-/-^* mice were strongly stained with alcian blue-PAS (Fig. 3a), it may be caused by an increased number of goblet cells. The colonic progenitor cells generated from Lgr5-expressing stem cells gradually differentiate into the two main epithelial lineages: absorptive and secretory lineage. The Hes1-expressing absorptive progenitors differentiate into all enterocytes, while Atoh1-expressing secretory progenitors differentiate into either enteroendocrine cells or goblet cells. We thus examined the mRNA expression of epithelial lineage markers, and found that the expression of *Lgr5* was decreased but that of *Atoh1* was increased in colon of *Gpr164^-/-^* mice (Fig. 4a). To further elucidate the role of GPR164 in intestinal epithelial differentiation, we performed genome-wide RNA sequencing by using colon samples from WT and *Gpr164^-/-^* mice. KEGG enrichment analysis revealed that the expression of genes related to the regulation of receptor activities were significantly changed in *Gpr164^-/-^*mice (Supplementary Fig. 3a). Notably, the expression of marker genes for enteroendocrine cells (*Chga*, *Gcg*, *Pyy*) and absorptive cells (*Slc26a2*, *Slc4a2*) were reduced, whereas these for goblet cells (*Reg4*, *Spink4*, *Fcgbp*) were elevated in *Gpr164^-/-^* mice (Fig. 4b). In addition, quantitative PCR analysis showed that the expression of *Tph-1* and *Vil1* was decreased but that of *Muc2* was increased in *Gpr164^-/-^*mice, confirming that a loss of *Gpr164* resulted in the enhanced differentiation into goblet cells (Fig. 4b). The proliferation and differentiation of colonic epithelial cells are regulated by Wnt/β-catenin signaling pathway. Upon activation of Frizzled and LRP5/6 receptors by binding Wnt ligands, β-catenin escapes from proteasomal degradation and accumulates in the nucleus, where it promotes transcription of genes involved in cell cycle progression by interacting with transcription factors of the T-cell factor/lymphoid enhancing factor (TCF/LEF) family (Fig. 4c). Therefore, Wnt signaling is essential for maintaining a proliferative phenotype of epithelial stem/progenitor cells in crypt, whereas the absence of this signal in the villus is important to allow epithelial cell differentiation. A heatmap of the gene expression profiles revealed that a large number of genes associated with Wnt signaling pathway were markedly changed, and ectopic expression of β-catenin in the villus was observed in *Gpr164^-/-^* mice (Fig. 4c, d). These results suggest that loss of *Gpr164* leads to ectopic expression of β-catenin in the villus, which causes abnormal proliferation (colonic hyperplasia shown in Fig. 2b-d) and differentiation (increased goblet cells and reduced enteroendocrine/absorptive cells as shown in Fig. 4b) of colonic epithelial cells. On the contrary, mRNA expression of *Lgr5* was reduced in *Gpr164^-/-^* mice, implying the possibility that colonic epithelial stem cells in the crypt cannot be received an appropriate Wnt signaling. It is well-known that dysregulation of Wnt signaling results in several diseases such as IBD and colorectal cancers^23,24^. Consistent with the data obtained in Fig. 2h and Fig. 3f, Kyoto Encyclopedia of Genes and Genome (KEGG) analysis showed that *Gpr164* deletion leads to disruption of epithelial barrier integrity and severe inflammation in the colon (Supplementary Fig. 3b, c). Therefore, our study indicates that loss-of-function mutation in *Gpr164* may play a pathogenic role in the development of IBD and cancers.

**Fig. 4.**
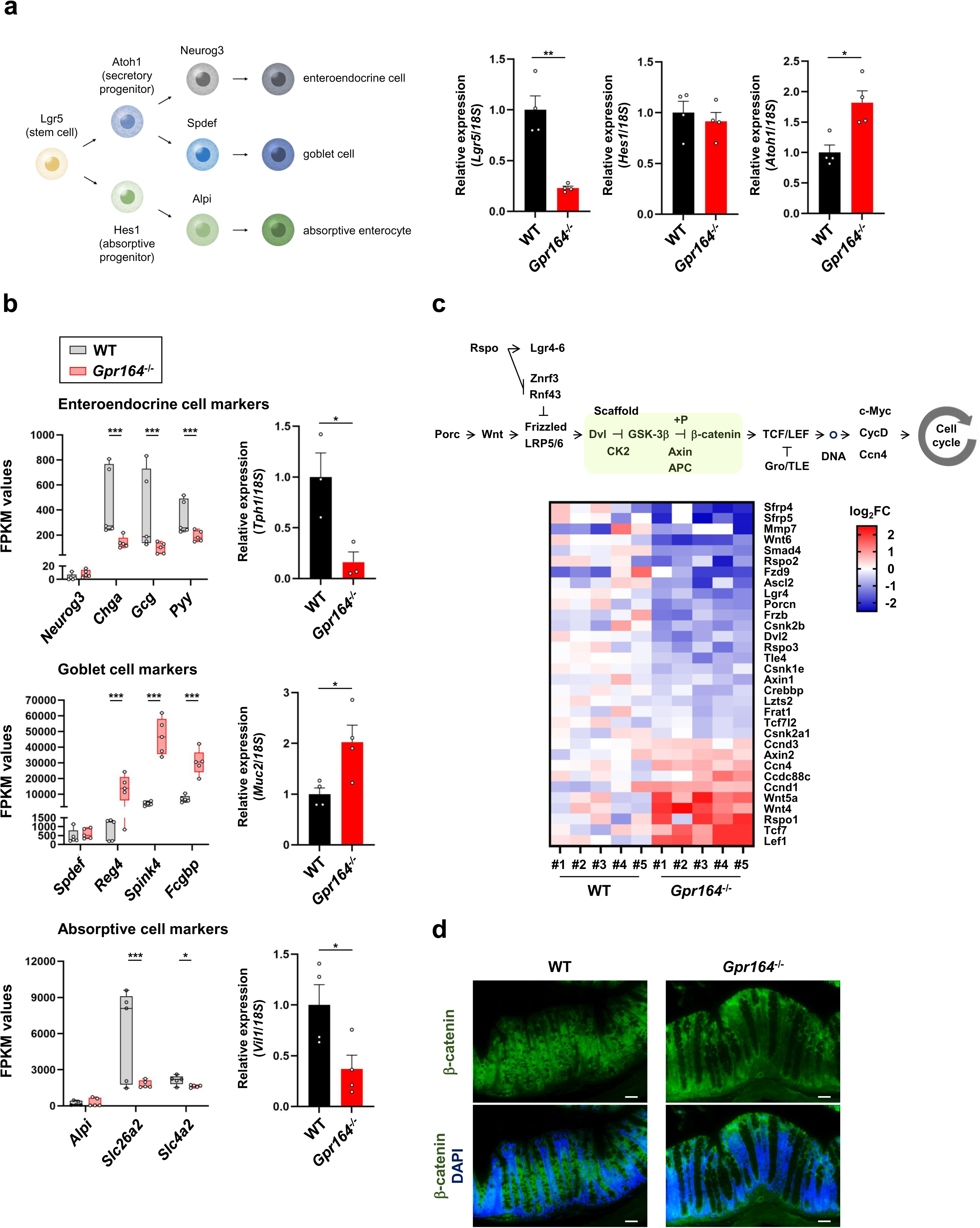
Aberrant Wnt signaling causes dysregulation of the colonic epithelial differentiation in *Gpr164*^-/-^ mice. (a) Schematic illustration of the intestinal epithelial lineage (left), and the mRNA expression levels of *Lgr5*, *Hes1* and *Atoh1* determined by qRT-PCR. Total RNA was extracted from colon tissue of WT and *Gpr164^-/-^*mice (n=4). Error bars represent the mean ±SEM. \**P* < 0.05, \*\**P* < 0.01 (Student’s *t*-test). (b) FPKM values of genes expressed in enteroendocrine cells (left, upper), goblet cells (left, middle) and absorptive cells (left, bottom). The mRNA expression levels of *Tph1* (right, upper), *Muc2* (right, middle) and *Vil1* (right, bottom) determined by qRT-PCR. Total RNA was extracted from colon tissue of WT and *Gpr164^-/-^* mice (n=3-4). Error bars represent the mean ±SEM. \**P* < 0.05, \*\*\**P* < 0.001 (Student’s *t*-test). (c) Schematic diagram of Wnt/β-catenin signaling pathway (upper), and heatmap of representative genes, involved in Wnt/β-catenin signaling, with significantly differences between WT and *Gpr164^-/-^* mice (bottom) (n=5). (d) Representative images of immunofluorescent staining for β-catenin. The cross-sections of colon obtained from WT and *Gpr164^-/-^* mice were stained with anti-β-catenin antibody (n=5). DAPI was used for nuclei staining. Scale bar, 100 μm.

### Inhibition of Wnt signaling ameliorates abnormalities in colon of *Gpr164*^-/-^ mice

To further elucidate the functional role of GPR164 in maintaining intestinal homeostasis, we examined the effect of Wnt inhibitor, PNU-74654, on the development of colonic epithelial abnormalities caused by *Gpr164* deletion. PNU-74654 acts as a Wnt/β-catenin antagonist by preventing TCF from binding to β-catenin, and inhibits tumor growth in a mouse model of colorectal cancer^25^. Intraperitoneal administration of PNU-74654 suppressed the hypertrophic phenotypes in the colon of *Gpr164^-/-^*mice (Fig. 5a, b). The anti-proliferative effects of PNU-74654 in *Gpr164^-/-^* mice were mediated through repressing increased expressions of *cyclin D1* and *c-Myc*, although decreased expression of *p21* and *p27* were partially restored (Fig. 5c and Supplementary Fig. 4). Also, a marked reduction in Ki67 protein expression of *Gpr164^-/-^* mice confirmed anti-proliferative effect of PNU-74654 (Fig. 5d), suggesting that accelerated proliferation in *Gpr164^-/-^* mice is caused by Wnt signaling overactivation. Next, we investigated the involvement of Wnt signaling in abnormal differentiation of colonic epithelial cells in *Gpr164^-/-^*mice. Upon treatment with PNU-74654, a lower and a higher expression level of *Vil1* and *Muc2*, respectively, in *Gpr164^-/-^* mice were restored to similar level in WT mice (Fig. 5e), and strong staining for mucin in the colon section of *Gpr164^-/-^* mice was attenuated (Fig. 5f). Therefore, these results suggest that aberrant Wnt signaling caused by *Gpr164* deletion leads to the pathophysiologic consequences such as abnormal differentiation in colonic epithelial cells. To examine whether dysregulation of Wnt signaling in *Gpr164^-/-^* mice affects the intestinal barrier function, mRNA expression of tight junction markers and plasma TNFα levels were determined. The reduced mRNA expression of *Occludin* and *Zo-1* and high levels of plasma TNFα were rescued to WT levels by administering PNU-74654 to *Gpr164^-/-^* mice (Fig. 5g, h). Together, GPR164 plays an important role in Wnt signaling-mediated colonic epithelial turnover, which is essential for maintaining the intestinal homeostasis such as barrier function.

**Fig. 5.**
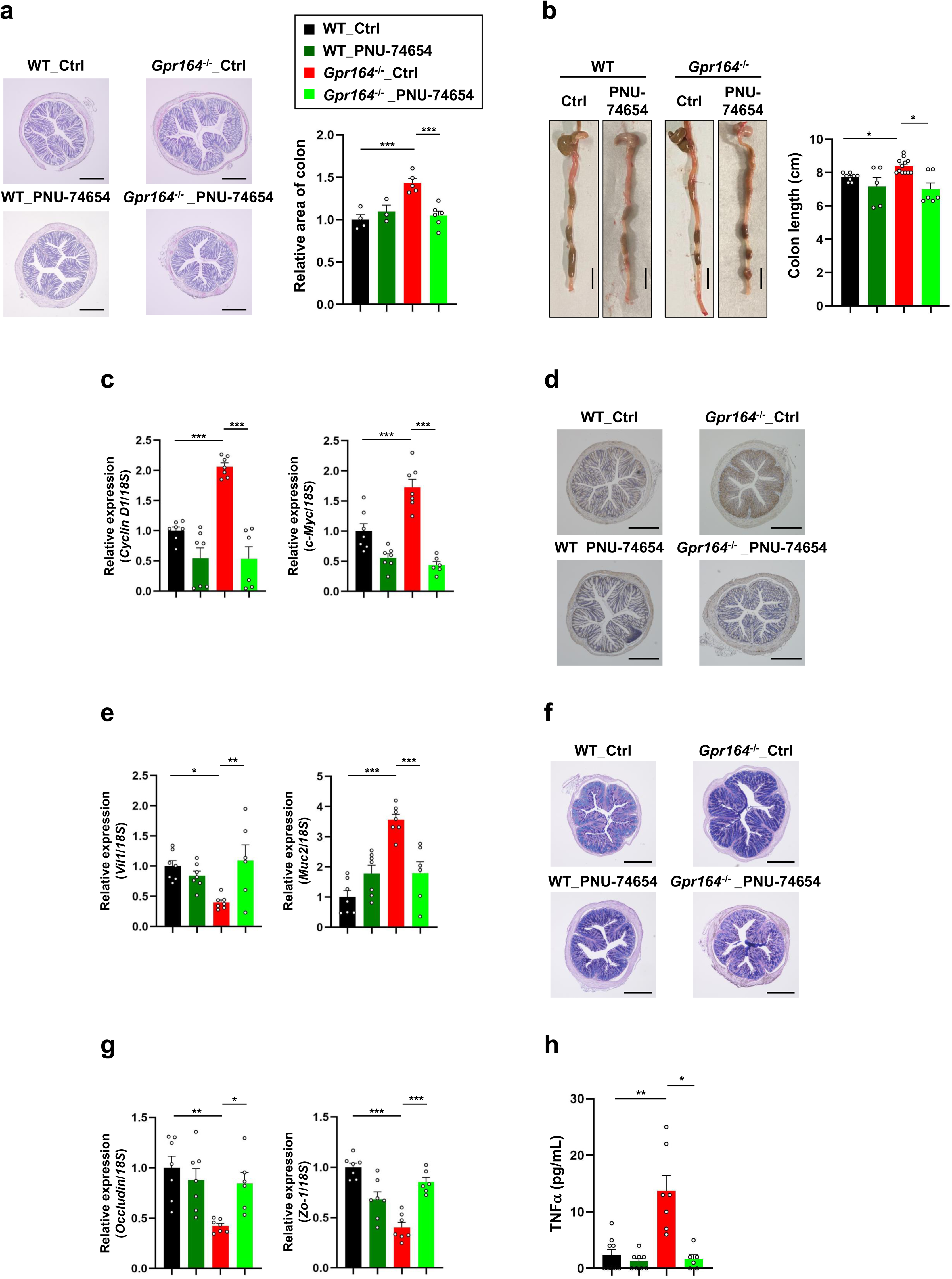
PNU-74654 ameliorates colonic abnormalities in *Gpr164*^-/-^ mice. (a) Representative images of colon sections stained with hematoxylin-eosin (HE). WT and *Gpr164^-/-^* mice were injected intraperitoneally with PNU-74654 (15 mg/kg body weight) every 2 days for 3 weeks. The cross-sections of colon obtained from WT and *Gpr164^-/-^* mice were stained with HE, and the area of colon was measured using ImageJ software (n=3-6). Scale bar, 500 μm. Error bars represent the mean ±SEM. \*\*\**P* < 0.001 (Tukey-Kramer test). (b) Representative images of colon (n=5-12). Scale bar, 1 cm. Error bars represent the mean ±SEM. \**P* < 0.05 (Tukey-Kramer test). (c) The mRNA expression levels of *Cyclin D1* and *c-Myc* determined by qRT-PCR. Total RNA was extracted from colon tissue of WT and *Gpr164^-/-^*mice (n=6-7). Error bars represent the mean ±SEM. \*\*\**P* < 0.001 (Tukey-Kramer test). (d) Representative images of immunohistochemical staining for Ki67 in colon. The cross-sections of colon obtained from WT and *Gpr164^-/-^* mice were stained with anti-Ki67 antibody. DAB was used for the detection of Ki67 protein, and hematoxylin was used for nuclei staining (n=3). Scale bar, 500 μm. (e) The mRNA expression levels of *Vil1* and *Muc2* determined by qRT-PCR. Total RNA was extracted from colon tissue of WT and *Gpr164^-/-^* mice (n=6-7). Error bars represent the mean ±SEM. \**P* < 0.05, \*\**P* < 0.01, \*\*\**P* < 0.001 (Tukey-Kramer test). (f) Representative images of colon sections stained with alcian blue-PAS (n=3). Scale bar, 500 μm. (g) The mRNA expression levels of *Occludin* and *Zo-1* determined by qRT-PCR. Total RNA was extracted from colon tissue of WT and *Gpr164^-/-^* mice (n=6-7). Error bars represent the mean ±SEM. \**P* < 0.05, \*\**P* < 0.01, \*\*\**P* < 0.001 (Tukey-Kramer test). (h) Plasma levels of TNFα (n=6-8). Error bars represent the mean ±SEM. \**P* < 0.05, \*\**P* < 0.01 (Dunn’s test).

### Butyrate enhances epithelial barrier function through GPR164 activation

The intestinal organoid is a 3D *in vitro* model with self-renewal capacity, and shows similar functionality to the tissue origin ^26^. Using colonic organoids, we assessed the role of ligand-activated GPR164 in the intestinal barrier maintenance. The colonic epithelial organoids isolated from *Gpr164^-/-^* mice showed no obvious differences compared to these from WT mice, when maintained in culture without passage (Fig. 6a). Interestingly, organoids from *Gpr164^-/-^* mice exhibited crypt-like budding structures after a single passage (Fig. 6a). The budding organoids possess similar features to the mature intestine^27^, thereby indicating a possibility that loss-of-function mutation in *Gpr164* is associated with accelerated differentiation and maturation of colonic epithelial cells. Consistent with *in vivo* findings, mRNA expression level of *Vil1* and *Occludin* was reduced but that of *Muc2* was increased in *Gpr164^-/-^* organoids (Fig. 6b). In addition, expressions of cytostatic genes, *p21* and *p27*, were decreased, although no significant changes in *Cyclin D1* and *c-Myc* mRNA expression were observed (Fig. 6b). Finally, we investigated the effect of butyrate on the intestinal barrier function using organoids. For the measurement of the organoid permeability, the luminal FITC-dextran which moves from the basolateral surface was examined^28^. FITC fluorescence in the lumen was detected in *Gpr164^-/-^* organoids treated with or without butyrate (Fig. 6c). Upon stimulation with palmitate, well-known as lipotoxic FFA^29^, increasing intensity of luminal FITC fluorescence was observed in both WT and *Gpr164^-/-^* organoid, and pretreatment with butyrate suppressed palmitate-induced increase in luminal FITC fluorescence in WT organoid but not in *Gpr164^-/-^* organoid (Fig. 6c). Palmitate has been reported to alter the expression of Hes1 and Muc2, thereby influencing the differentiation of certain types of cells such as goblet cells^30^. Therefore, besides pro-inflammatory property^29^, palmitate induces lipotoxicity through impairment of epithelial differentiation. Taken together, our results indicate that butyrate-mediated activation of GPR164 plays a crucial role in protecting the barrier function in part through regulating proliferation and differentiation of colonic epithelial cells.

**Fig. 6.**
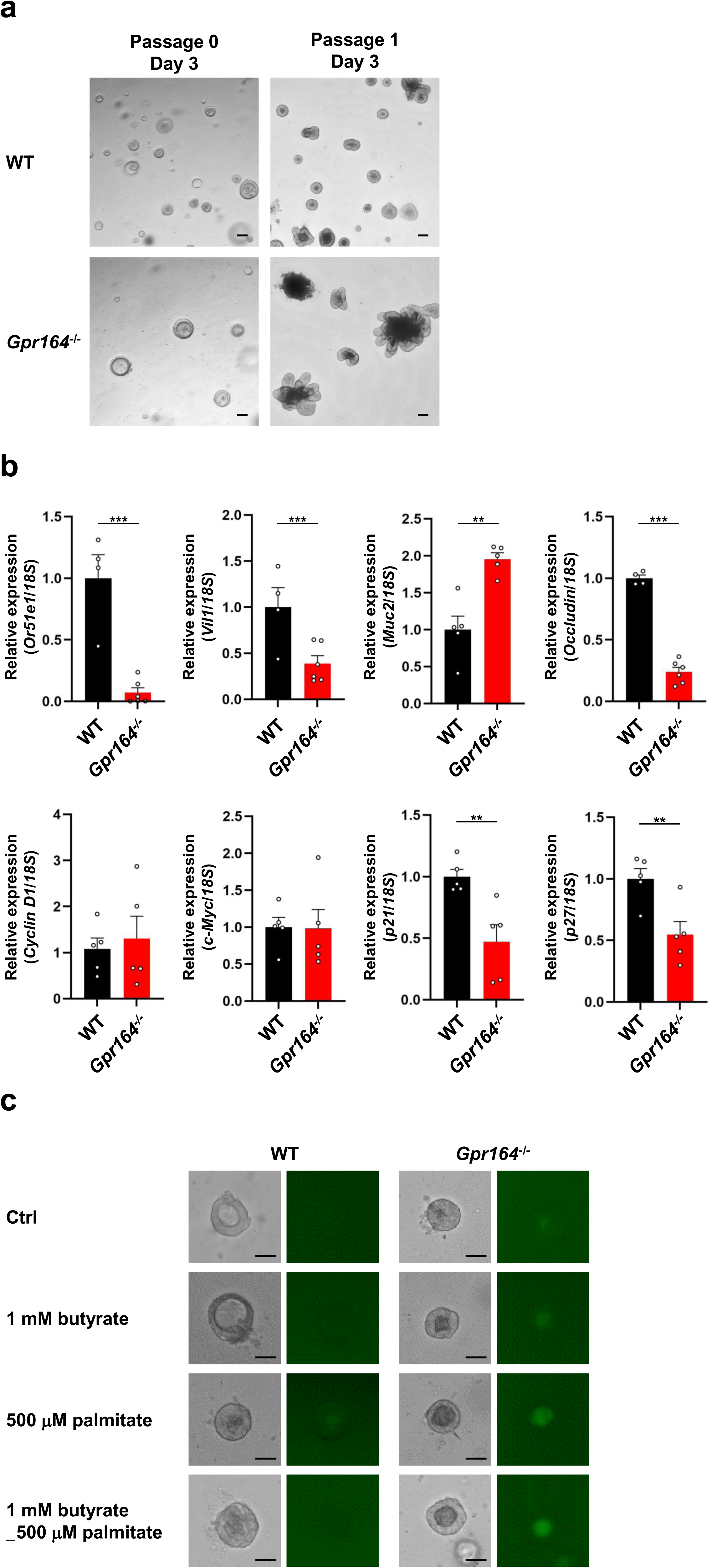
Butyrate-mediated enhancement of epithelial barrier function is lost in *Gpr164*^-/-^ mice. (a) Representative images of colonic organoids isolated from WT and *Gpr164^-/-^*mice (n=3). Scale bar, 100 μm. (b) The mRNA expressions in colonic organoids determined by qRT-PCR. Total RNA was extracted from colonic organoids isolated from WT and *Gpr164^-/-^* mice (n=4-6). Error bars represent the mean ±SEM. \*\**P* < 0.01, \*\*\**P* < 0.001 (Student’s *t*-test). (c) Representative images of fluorescence in colonic organoids. Colonic organoids isolated from WT and *Gpr164^-/-^* mice were preincubated for 24 h in the presence or absence of 1 mM sodium butyrate, and followed by stimulation with or without 500 μM sodium palmitate for 24 h. Leakage of FITC-dextran into the lumen was assessed by fluorescence (n=3). Scale bar, 50 μm.

## Discussion

In the present study, we revealed the pivotal role of a novel FFAR, GPR164, on the intestinal barrier function using *Gpr164^-/-^* mice. To date, there have been no reports describing *in vivo* functions of GPR164 by using *Gpr164^-/-^* mice. Thus, our study provides the first line of evidence how GPR164 maintains the physiological homeostasis. A loss-of-function mutation in *Gpr164* resulted in a disruption of the intestinal barrier function (Fig. 2h), which was caused by aberrant Wnt signaling because expression of β-catenin was observed in the intestinal villus tips (Fig. 4d). This ectopic β-catenin expression promoted proliferation of colonic epithelial cells, and altered the epithelial lineage. As a consequence, the colon of *Gpr164^-/-^* mice had an enlarged volume, and epithelial cells preferentially differentiated into goblet cells rather than into enteroendocrine cells or absorptive cells (Fig. 4a, b). However, thickness of mucus layer was reduced in colon of *Gpr164^-/-^* mice (Fig. 3b), implying the possibility that accumulated mucins could not be released from goblet cells. Butyrate is an important gut microbial metabolite for the maintenance of mucus barrier function through promoting expression and release of mucins^31,32^. Considering our results shown in Fig. 3g, decreased level of butyrate may be insufficient to stimulate mucin secretion. Therefore, further studies are required to understand the molecular mechanisms underlying mucus barrier function, including a possibility that butyrate-mediated activation of GPR164 facilitates mucin secretion. Besides the mucus barrier, GPR164 plays an important role in the tight junction barrier. *Gpr164^-/-^* mice showed significant reduced expression of absorptive/enteroendocrine cell markers and tight junction markers (Fig. 2g and Fig. 3b). These reduced expressions were restored to WT levels by the treatment with Wnt signaling inhibitor, PNU-74654 (Fig. 5e, g), suggesting that colonic epithelial cells of *Gpr164^-/-^* mice may have defects in the formation of organized monolayer because of the abnormal Wnt signaling. Furthermore, *OR51E1* KO Caco-2 cells also showed hyperproliferation and intestinal barrier dysfunction (Fig. 1d, f), providing evidence for an essential role of GPR164 in maintaining the intestinal barrier function both *in vivo* and *in vitro* models. Together, these results suggest that aberrant activation of Wnt signaling caused by *Gpr164* absent leads to dysfunction of mucus barrier and tight junction barrier.

Defects in the intestinal barrier function are strongly associated with gut microbial dysbiosis. Indeed, we found the marked alterations in microbial composition and increased F/B ratio in *Gpr164^-/-^*mice (Fig. 3c-e). A previous report has shown that changes in the luminal microenvironment caused by pathogen challenge and dietary manipulations affect the *Gpr164* gene expression^4^, indicating a positive correlation between GPR164 and gut microbial diversity. Both the intestinal barrier dysfunction and gut microbial dysbiosis are the characteristic features of IBD, and the chronic inflammation in IBD patients increases a risk of developing colorectal carcinoma (CRC)^33^. Here we showed that *Gpr164^-/-^* mice exhibited severe colonic inflammation and Wnt signaling abnormalities (Fig. 3f, Fig. 4 and Fig. 5). Overactivation of Wnt signaling is also known as a hallmark of CRC, and therefore, our findings highlight GPR164 as an attractive therapeutic target for IBD and CRC. Overall, this study uncovers the physiological functions of GPR164, which will help us understand the molecular links between diet and physiology.

## Acknowledgments

This study was supported by research grants from the JSPS KAKENHI (22K21208 and 23K16794 to T.I.), Takeda Science Foundation (to T.I.) and JST-MOONSHOT (JPMJMS2023 to I.K.).

## Author contributions

T.I. performed the experiments and wrote the paper. Y.M. performed the experiments and interpreted the data. K.W. performed the experiments and interpreted the data. A.N. performed the experiments. M.Y. performed the experiments. M.I. performed the experiments. N.S. performed the experiments. H.K. performed the experiments. I.K. supervised the projects and interpreted the data. T.I. and I.K. had primary responsibility for the final content. All authors read and approved the final version of the manuscript.

## Methods

### Animals

Male C57BL/6J mice were purchased from Japan SLC and *Or51e1* gene knockout (*Gpr164^-/-^*) mice were generated by using the CRISPR/Cas9 system in wild-type C57BL/6J zygotes (Supplementary Fig. 2). C57BL/6J and *Gpr164^-/-^* mice were housed at a temperature of 24 °C and 50% relative humidity under a 12 h light/dark cycle. For high-fat diet (HFD) feeding study, 6-week-old male mice were fed with either a normal chow diet (ND) (CE-2; CLEA Japan) or a HFD (D12492, 60% kcal fat; Research diet) for 10 weeks. For the assessment of intestinal permeability, mice withdrawn from food and water for 4 h were administrated a freshly prepared FITC-dextran (4 kDa; Sigma) at 0.6 g/kg body weight by oral gavage. After 3 h, blood was collected from the inferior vena cava, and plasma was separated by immediate centrifugation at 7000g for 5 min at 4 °C. The fluorescent intensity of FITC-dextran within plasma samples was measured by FlexStation3 fluorescence plate-reader (emission 520 nm, excitation 490 nm). A standard curve was generated by serial dilution of FITC-dextran, and used for determining concentration of FITC-dextran in plasma samples. All animal experiments were performed in accordance with the guidelines of the Committee in the Ethics of Animal Experiments of Kyoto University Animal Experimental Committee (Lif-K24002). All efforts were made to minimize suffering.

### Cell culture

Caco-2 and HEK293 cells were maintained in Dulbecco’s modified Eagle’s medium (DMEM) supplemented with 10% fetal calf serum, penicillin and streptomycin at 37°C in 5% CO_2_. To generate Or51e1-overexpressing cells, HEK293 cells were co-transfected with pcDNA3.1_HA-*Or51e1* and pcDNA3.1_*Rtp1S* (or *RTP4*) using lipofectamine 2000 (Invitrogen). *OR51E1*-deficient (*OR51E1* KO) Caco-2 cells were generated using the CRISPR/Cas9 system. Single guide RNA (sgRNA) targeting *OR51E1* (5’-atccgggtcaatgtcgtcta-3’) was designed by using the online software CRISPOR (http://crispor.tefor.net). The sgRNA was cloned into peSpCAS9(1.1)-2xsgRNA vector (Addgene, #80768), and the recombinant peSpCAS9(1.1)-2xsgRNA and pDonor-tBFPNLS-Neo (Addgene, #80766) were co-transfected into Caco-2 cells using Lipofectamine 2000 (Invitrogen). For the selection of *OR51E1* KO cells, transfected cells were maintained in medium containing 250 mg/mL G418 (FUJIFILM Wako), and the colony grown from a single cell was isolated. To evaluate the intestinal barrier function, Caco-2 cells were seeded on a cell culture insert (0.4 μm pore; Falcon) and transepithelial electrical resistance (TER) value was measured using Millicell-ERS system (Millipore).

### Quantitative RT-PCR

Total RNA was extracted using an RNAiso Plus reagent (TAKARA), and cDNA was synthesized by using Moloney murine leukemia virus reverse transcriptase (Invitrogen). Quantitative PCR analysis was performed using the StepOne real-time PCR system (Applied Biosystems) with SYBR Premix Ex Taq II (TAKARA). Each value was normalized to *18S* rRNA and calculated using the 2-ΔΔCt method. The primer sequences are shown in Supplementary Table 1.

### cAMP measurement

HEK293 cells transfected with either empty or Or51e1 expression vector were maintained in serum-free DMEM containing 500 μM of 3-isobutyl 1-methylxantine (Sigma) for 30 min. Cell were then stimulated with individual FFAs at 1 mM concentration for 10 min. The cAMP levels were determined using cAMP EIA kit (Cayman Chemical) according to the manufacturer’s instructions. Samples treated with 2 μM folskolin (Sigma) were used as positive controls.

### Crystal violet staining

Caco-2 cells were seeded in 24-well plate (1×10^5^ cells/well) and fixed with 4% paraformaldehyde at the indicated times. Cells were then stained with 0.1% crystal violet (FUJIFILM Wako) for 15 min, and washed with PBS three times. The dye was dissolved in 95% ethanol, and absorbance was measured at OD_590_ nm.

### Immunoblot and immunofluorescence analysis

Immunoblotting was performed according to a standard protocol using anti-p53 antibody (Santa Cruz, sc-126) and anti-tubulin antibody (Sigma, T5168). α-Tubulin was used as a loading control. For immunofluorescence analysis, 10% formalin-fixed colon sample was embedded in paraffin and cross-sectioned at 4 μm thickness (Kyoto Institute of Nutrition & Pathology, Inc). The cross-section was labeled with anti-β-catenin primary antibody (Proteintech, 51067-2-AP) followed by staining with Alexa Fluor 488, and observed with a fluorescence microscope (Keyence, BZ-X710). DAPI was used for nuclei staining.

### HE staining and immunohistochemistry

Hematoxylin and eosin (HE) staining was performed by a standard method. The paraffin-embedded cross-section was deparaffinized and stained with eosin and hematoxylin. Immunohistochemistry was performed according to a standard protocol. Briefly, the paraffin-embedded cross-section was deparaffinized, and incubated with blocking serum after antigen retrieval. The anti-Ki67 antibody (Proteintech, 28074-1-AP) was used as a primary antibody, and peroxidase-conjugated secondary antibody (Vector Laboratories, PI-1000) was used. The cross-section was visualized with diaminobenzidine (DAB) and counterstained with hematoxylin.

### Alcian blue-periodic acid Schiff (PAS) staining

The colon sample was fixed in Carnoy’s solution (FUJIFILM Wako) and embedded in paraffin. Alcian blue-PAS staining of mucins was performed using alcian blue-PAS stain kit (Scy Tek), following the manufacturer’s instructions. In short, deparaffinized colon section was treated with 3% acetic acid for 2 min, followed by staining with alcian blue (pH 2.5) for 15 min. After washing with running tap water, the tissue section was treated with periodic acid for 5 min. The tissue section was then incubated with schiff’s solution, and stained with hematoxylin.

### Measurement of SCFAs

SCFAs in the colon contents were determined as described previously^34^. In brief, the samples containing internal control (2-ethyl butyrate) were mixed with diethyl ether and centrifuged at 3000 × g for 5 min. SCFA-containing ether layers were collected and subjected to gas chromatography-mass spectrometry using a GCMS-QP2010 Ultra system (Shimadzu). Using the calibration curves for SCFAs, SCFA concentration in each sample was determined.

### 16S rRNA sequencing of gut microbes

Fecal DNA was extracted using FastDNA SPIN kit for feces (MP Biomedicals), according to the manufacturer’s instructions. To determine the microbial taxa, the V3-V4 regions of the bacterial 16S rRNA gene were amplified using following primers; (forward) 5’-TCGTCGGCAGCGTCAGATGTGTATAAGAGACAGCCTACGGGNGG CWGCAG-3’, (reverse) 5′-GTCTCGTGGGCTCGGAGATGTGTATAAGAGACAGGA CTACHVGGGTATCTAATCC-3. Amplified products from each sample were purified by AMPure XP (Beckman Coulter) and sequenced using the MiSeq platform (Illumina). Raw data were processed using quantitative insights into microbial ecology (QIIME) pipeline, and analyzed by the MiSeq reporter software with the SILVA database (Illumina). The diversity of gut microbiota was determined by the QIIME script core_diversity_analyses.py. The statistical significance of difference between groups was assessed by a permutational multivariate analysis of variance (QIIME script compare_categories.py).

### RNA sequencing

Total RNA was extracted from the colon of WT and *GPR164^-/-^*mice using an RNAiso Plus reagent (TAKARA) and RNeasy mini kit (Qiagen). The quality of RNA samples was determined by Agilent 2100 Bioanalyzer with an RNA 6000 Nano kit (Agilent Technologies). RNA sequencing libraries were created using the NEBNext® Ultra™ II Directional RNA Library Prep Kit (Illumina) and NEBNext Multiplex Oligos for Illumina (Dual Index Primers Set 1). Paired-end sequencing was performed using an Illumina NovaSeq 6000 (150 bp read lengths; approximately 4 Gb). The data obtained from RNA sequencing were trimmed with trimmomatic (v0.39) software to eliminate adapters and inadequate quality bases^35^, and FastQC (v0.11.8.-2) software was used for quality control of the trimmed sequence. The reads were aligned to the mouse reference genome (NCBI GRCm39) using STAR software (v2.7.10a). To obtain differentially expressed genes (DEGs) across all comparisons, the raw read counts were subjected to relative log expression normalization. Based on the following criteria, DEGs were determined; a false discovery rate (FDR)-adjusted p-value < 0.05 (Benjamini-Hochberg procedure) and an absolute log2 fold change > 0.5. Gene Set Enrichment Analysis was performed using the Kyoto Encyclopedia of Genes and Genomes (KEGG) database (http://www.genome.jp/kegg/).

### Organoid culture

Mouse colonic segments of WT and *GPR164^-/-^* mice were collected in a PBS solution containing 5 mM EDTA, and rotated for 40 min at 4°C. The colonic crypts were isolated from the segments by repetitive pipetting, and filtered through a cell strainer. The isolated crypts were embedded in Matrigel, and maintained in human IntestiCult organoid growth medium (Veritas) containing penicillin and streptomycin. Organoids were maintained at 37°C in 5% CO_2_, and the medium was changed every other day. The assessment of barrier function using organoids was performed as described previously^28^. Briefly, the colonic organoids were preincubated with or without 1 mM sodium butyrate for 24 h, and stimulated with either 500 μM sodium palmitate or control DMSO for 24 h. After stimulation, the organoids were washed with PBS, and incubated with FITC-dextran (4kDa; Sigma) at 1.25 μM concentration for 1 h. To remove FITC-dextran from the media, organoids were gently washed with PBS, and fluorescence within the organoids was observed with a fluorescence microscope (Keyence, BZ-X710).

### Statistical analysis

All values are shown as mean ± standard error of the mean (SEM). Statistical analyses were performed using GraphPad Prism software (GraphPad Software). The Shapiro-Wilk test was used for the assessment of data normality. The statistical significance of differences between two groups was determined by a two-tailed unpaired Student’s *t*-test, while that of differences among multiple groups was determined by one-way ANOVA followed by the Turkey-Krammer test or Dunn’s test. Statistical significance was defined as P < 0.05.

### Data availability

The source data shown in RNA-seq analysis and 16S rRNA sequence have been deposited into the DNA Data Bank of Japan (DDBJ) under the accession no. E-GEAD-1039 and DRA020128, respectively. The source data underlying Figs. 1-6 and Supplementary Figs. 1,2,4 have been deposited into the Dryad repository (https://doi.org/10.5061/dryad.ghx3ffc0k).

## References

1. Bellono, N.W., et al. Enterochromaffin cells are gut chemosensors that couple to sensory neural pathways. Cell 170, 185–198 (2017).

2. Saito, H., Chi, Q., Zhuang, H., Matsunami, H. & Mainland, J.D. Odor coding by a mammalian receptor repertoire. Sci. Signal. 2, ra9 (2009).

3. Halperin Kuhns, V.L., et al. Characterizing novel olfactory receptors expressed in the murine renal cortex. Am. J. Physiol. Renal Physiol. 317, F172–F186 (2019).

4. Priori, D., et al. The olfactory receptor OR51E1 is present along the gastrointestinal tract of pigs, co-localizes with enteroendocrine cells and is modulated by intestinal microbiota. PLoS One 10, e0129501 (2015).

5. Han, Y.E., et al. Olfactory receptor OR51E1 mediates GLP-1 secretion in human and rodent enteroendocrine L cells. J. Endocr. Soc. 2, 1251–1258 (2018).

6. Shimizu, H, et al. Dietary short-chain fatty acids intake improves the hepatic metabolic 384 condition via FFAR3. Sci. Rep. 9, 16574 (2019).

7. Ikeda, T., Nishida, A., Yamano, M. & Kimura, I. Short-chain fatty acid receptors and gut microbiota as therapeutic targets in metabolic, immune, and neurological diseases. Pharmacol. Ther. 239, 108273 (2022).

8. Topping, D.L. & Clifton, P.M. Short-chain fatty acids and human colonic function: Roles of resistant starch and nonstarch polysaccharides. Physiol. Rev. 81, 1031–1064 (2001).

9. Mowat, A.M. & Agace, W.W. Regional specialization within the intestinal immune system. Nat. Rev. Immunol. 14, 667–685 (2014).

10. Furusawa, Y., et al. Commensal microbe-derived butyrate induces the differentiation of colonic regulatory T cells. Nature 504, 446–450 (2013).

11. Vernia, P., et al. Short-chain fatty acid topical treatment in distal ulcerative colitis. Aliment. Pharmacol. Ther. 9, 309–313 (1995).

12. Peng, L., Li, Z.R., Green, R.S., Holzman, I.R., & Lin, J. Butyrate enhances the intestinal barrier by facilitating tight junction assembly via activation of AMP-activated protein kinase in Caco-2 cell monolayers. J. Nutr. 139, 1619–1625 (2009).

13. Willemsen, L.E., Koetsier, M.A., van Deventer, S.J., & van Tol, E.A. Short chain fatty acids stimulate epithelial mucin 2 expression through differential effects on prostaglandin E1 and E2 production by intestinal myofibroblasts. Gut 52,1442–1447 (2003).

14. Pint, D. & Clevers, H. Wnt control of stem cells and differentiation in the intestinal epithelium. Exp. Cell Res. 306, 357–363 (2005).

15. Weng, J., Wang, J., Hu, X., Wang, F., Ittmann, M. & Liu, M. PSGR2, a novel G-protein coupled receptor, is overexpressed in human prostate cancer. Int. J. Cancer 118, 1471–1480 (2006).

16. Maßberg, D., et al. The activation of OR51E1 causes growth suppression of human prostate cancer cells. Oncotarget 7, 48231–48249 (2016).

17. Giandomenico, V., Cui, T., Grimelius, L., Öberg, K., Pelosi, G. & Tsolakis, A.V. Olfactory receptor 51E1 as a novel target for diagnosis in somatostatin receptor-negative lung carcinoids. J. Mol. Endocrinol. 51, 277–286 (2013).

18. Cui, T., et al. Olfactory receptor 51E1 protein as a potential novel tissue biomarker for small intestine neuroendocrine carcinomas. Eur. J. Endocrinol. 168, 253–261 (2013).

19. Zhuang, H. & Matsunsmi, H. Synergism of accessory factors in functional expression of mammalian odorant receptors. J. Biol. Chem. 282, 15284–15293 (2007).

20. Pronin, A. & Slepak, V. Ectopically expressed olfactory receptors OR51E1 and OR51E2 suppress proliferation and promote cell death in a prostate cancer cell line. J. Biol. Chem. 296, 100475 (2021).

21. Johansson, E.V.M, Holmén Larsson, M. J. & Hansson, C.G. The two mucus layers of colon are organized by the MUC2 mucin, whereas the outer layer is a legislator of host-microbial interactions. Proc. Natl. Acad. Sci. USA. 108, 4659–4665 (2010).

22. Swidsinski, A., et al. Comparative study of the intestinal mucus barrier in normal and inflamed colon. Gut 56, 343–350 (2007).

23. Aust, D.E., et al. The APC/beta-catenin pathway in ulcerative colitis-related colorectal carcinomas: a mutational analysis. Cancer 94, 1421–1427 (2002).

24. Sparks, A.B., Morin, P.J., Vogelstein, B. & Kinzler, K.W. Mutational analysis of the APC/beta-catenin/Tcf pathway in colorectal cancer. Cancer Res. 58, 1130–1134 (1998).

25. Amerizadeh, F., et al. Inhibition of the Wnt/β-catenin pathway using PNU-74654 reduces tumor growth in in vitro and in vivo models of colorectal cancer. Tissue Cell 77, 101853 (2022).

26. Sasaki, N. & Clevers, H. Studying cellular heterogeneity and drug sensitivity in colorectal cancer using organoid technology. Curr. Opin. Genet. Dev. 52, 117–122 (2018).

27. Onozato, T., et al. Generation of budding-like intestinal organoids from human induced pluripotent stem cells. J. Pharm. Sci. 110, 2637–2650 (2021).

28. Hu, X., et al. Aryl hydrocarbon receptor utilises cellular zinc signals to maintain the gut epithelial barrier. Nat. Commun. 14, 5431 (2023).

29. Ohue-Kitano, R., et al. Medium-chain fatty acids suppress lipotoxicity-induced hepatic fibrosis via the immunomodulating receptor GPR84. JCI Insight. 8, e165469 (2023).

30. Fillippello, A., et al. Molecular effects of chronic exposure to palmitate in intestinal organoids: A new model to study obesity and diabetes. Int. J. Mol. Sci. 23, 7751 (2022).

31. Willemsen, L.E.M., Koetsier, M.A.K., Deventer, S.J.H. & Tol E.A.F. Short chain fatty acids stimulate epithelial mucin 2 expression through differential effects on prostaglandin E_1_ and E_2_ production by intestinal myofibroblasts. Gut. 52, 1442–1447 (2003).

32. Jung, T.H., Han, K.S., Park, J.H. & Hwang, H.J. Butyrate modulates mucin secretion and bacterial adherence in LoVo cells via MAPK signaling. PLoS One. 17, e0269872 (2022).

33. Brackmann, S., et al. Relationship between clinical parameters and the colitis-colorectal cancer interval in a cohort of patients with colorectal cancer in inflammatory bowel disease. Scand. J. Gastroenterol. 44, 46–55 (2009).

34. Miyamoto, J., et al. Ketone body receptor GPR43 regulates lipid metabolism under ketogenic conditions. Proc. Natl. Acad. Sci. USA. 116, 23813–23821 (2019).

35. Bolger, A.M., Lohse, M. & Usadel, B. Trimmomatic: a flexible trimmer for Illumina sequence data. Bioinformatics 30, 2114–2120 (2014).

