## Supplementary information for "The pivotal role of a novel free fatty acid receptor GPR164 in the intestinal barrier function"

**Supplementary Fig. 1 | Generation of Or51e1-overexpressing and *OR51E1*-knockout cells.**

(a) Representative images of immunofluorescent staining for mouse Or51e1 in HEK293 cells. Cells were co-transfected with HA-tagged Or51e1 and receptor-transporting protein (left; RTP1S, right; RTP4), and stained with anti-HA antibody (green). DAPI was used for nuclei staining. Scale bar, 50 μm. (b) *OR51E1* expression in Caco-2 cells. *OR51E1*-deficient (*OR51E1* KO) cells were generated using the CRISPR/Cas9 system. For the detection of *OR51E1* expression, PCR amplification was done with the indicated primers, and the PCR products were separated on 1% agarose gels (left). The *OR51E1* mRNA expression level was determined by qRT-PCR (n=3) (right). Error bars represent the mean ±SEM. $^{*}P < 0.05$ (Student's *t*-test). (c) Representative image of p53 protein expression. Cell extracts from WT or *OR51E1* KO cells were subjected to immunoblot analysis using an anti-p53 or anti-αTubulin antibody.

**Supplementary Fig. 2 | Generation of *Or51e1*-knockout (*Gpr164*$^{-/-}$) mice.**

(a) Schematic representation of CRISPR/Cas9 targeting sites in *Or51e1* gene. *Or51e1* gene knockout (*Gpr164*$^{-/-}$) mice were generated by using the CRISPR/Cas9 system in wild-type C57BL/6J zygotes. Bold letters indicate the coding region of *Or51e1* gene. Red or Green letters indicate guide RNA (gRNA) and protospacer adjacent motif (PAM), respectively. (b) For the detection of the wild-type and mutant alleles, PCR amplification was done with the indicated primers, and the PCR products were separated on 1% agarose gels. (c) The *Or51e1* mRNA expression level in colon was determined by qRT-PCR (n=3). Error bars represent the mean ±SEM. $^{**}P < 0.01$ (Student's *t*-test).

**Supplementary Fig. 3 | Genome-wide RNA sequencing of *Gpr164*$^{-/-}$ mice.**

(a) KEGG enrichment analysis involved in the molecular function in colon of *Gpr164*$^{-/-}$ mice (n=5). P values were adjusted based on the false discovery rate (FDR). (b, c) KEGG pathway enrichment related to tight junction (b) and inflammatory bowel disease (c). Increased or decreased levels of gene expressions are shown in red or blue, respectively.

**Supplementary Fig. 4 | Cell cycle gene expressions in PNU-74654-treated *Gpr164*$^{-/-}$ mice.**

The mRNA expression levels of cell cycle genes determined by qRT-PCR (n=6-7). WT and *Gpr164*$^{-/-}$ mice were injected intraperitoneally with PNU-74654 (15 mg/kg body weight, every 2 days for 3 weeks), and total RNA was extracted from colon tissue of WT and *Gpr164*$^{-/-}$ mice (n=6-7). Error bars represent the mean ±SEM. **P* < 0.05, ***P* < 0.01 (Tukey-Kramer test; right, Dunn's test; left,).

**Supplementary Table 1 | Primer sequences used in this study.**

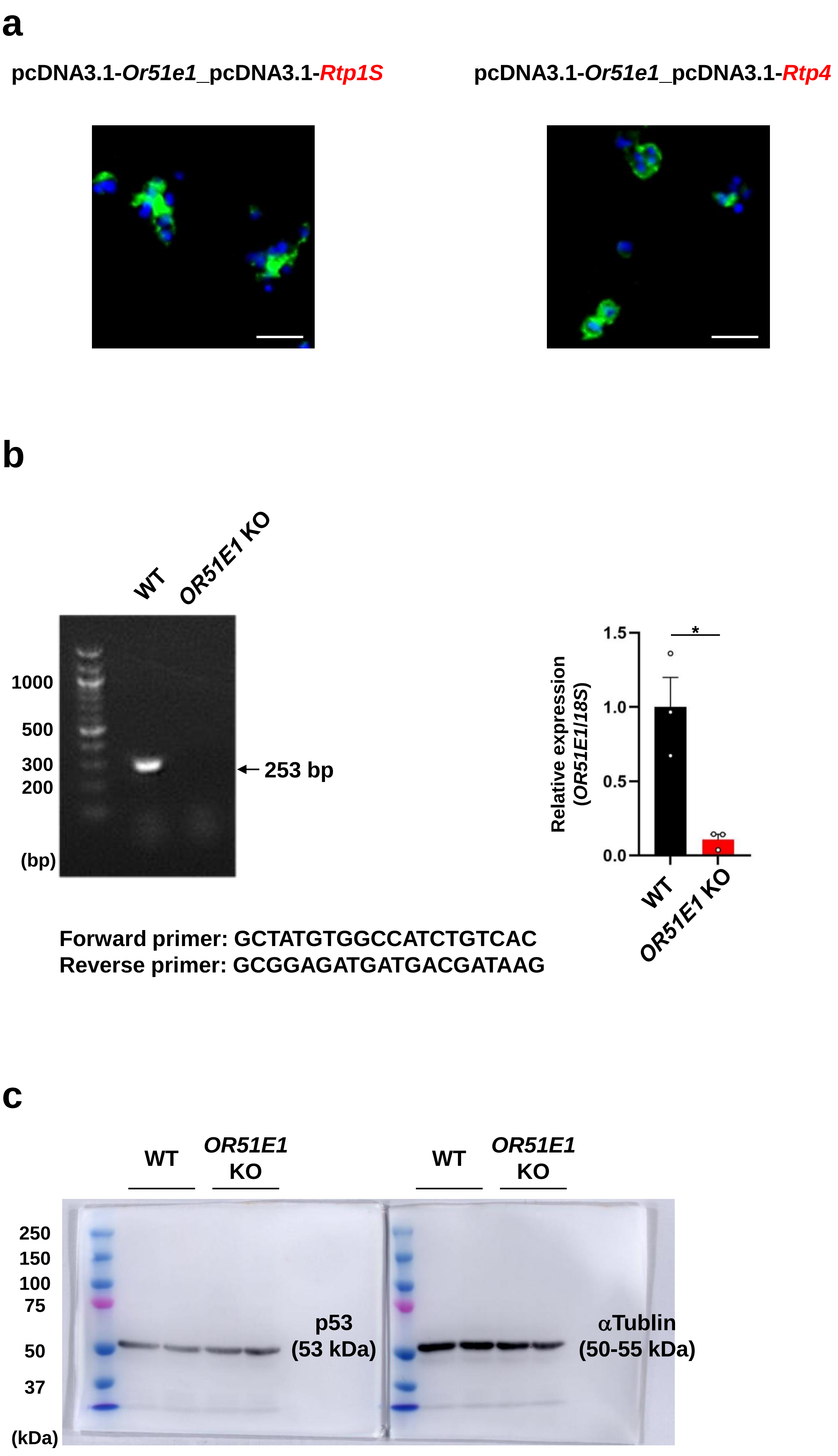


Supplementary Figure 1

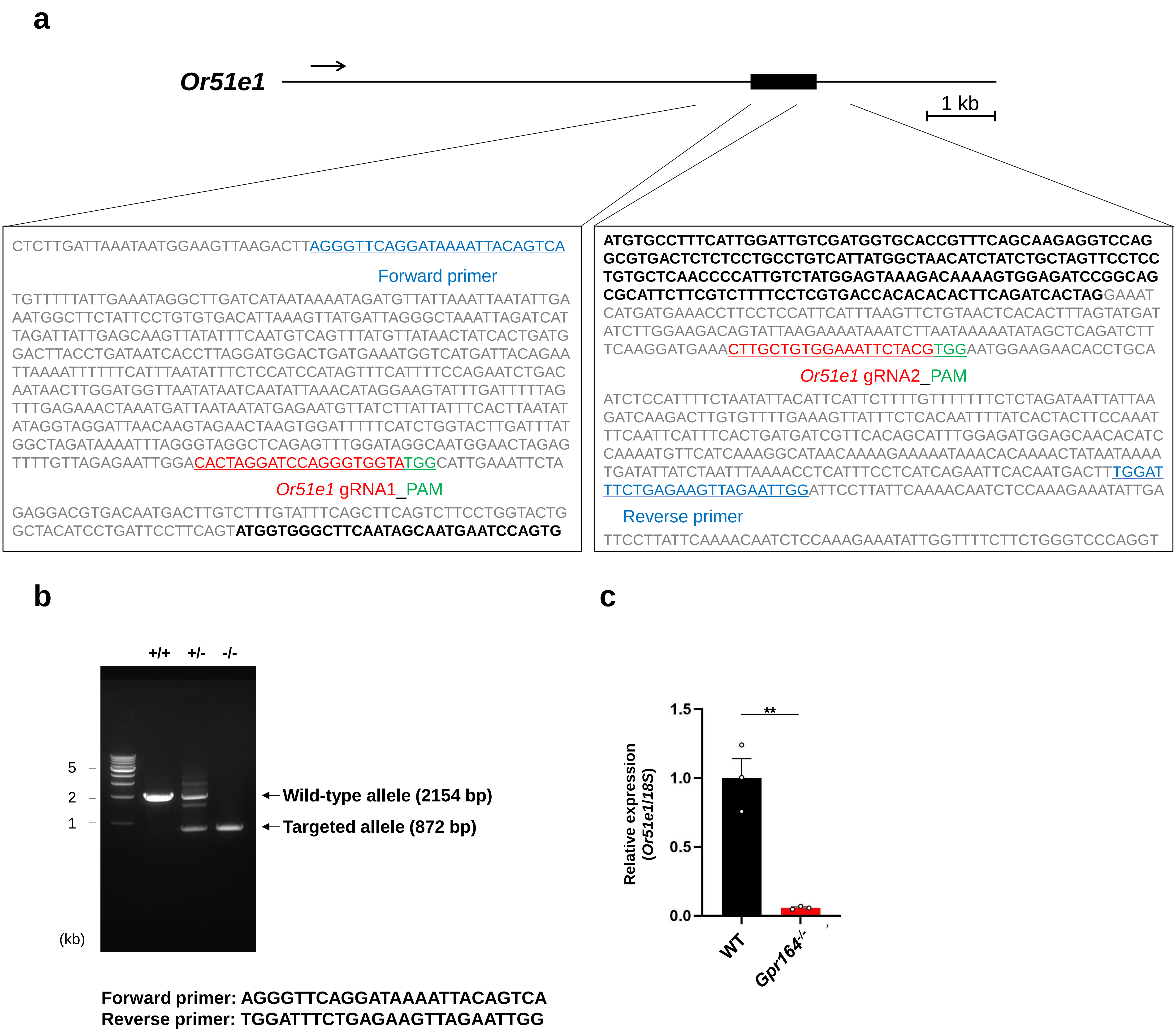


**Supplementary Figure 2**

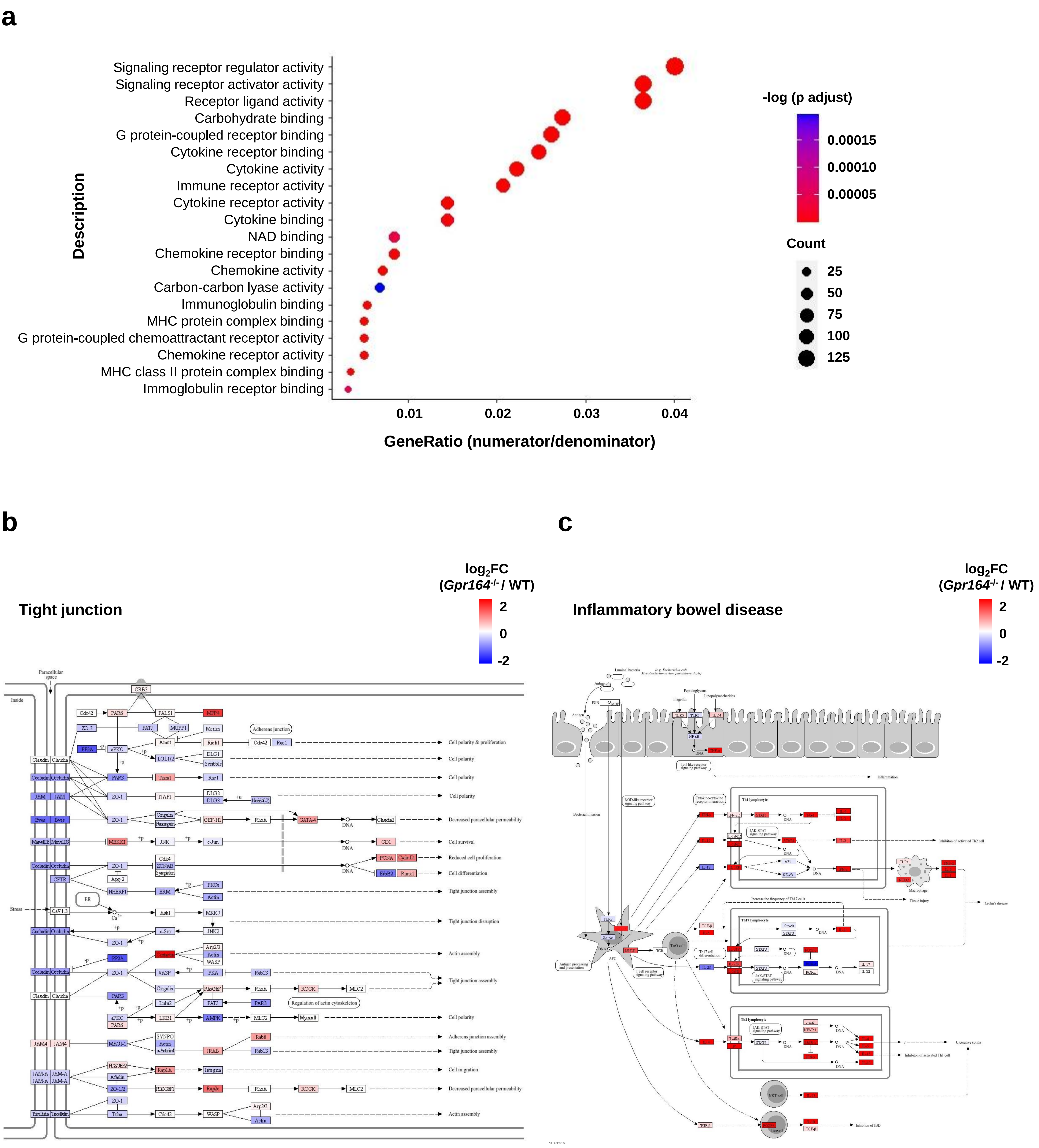


**Supplementary Figure 3**

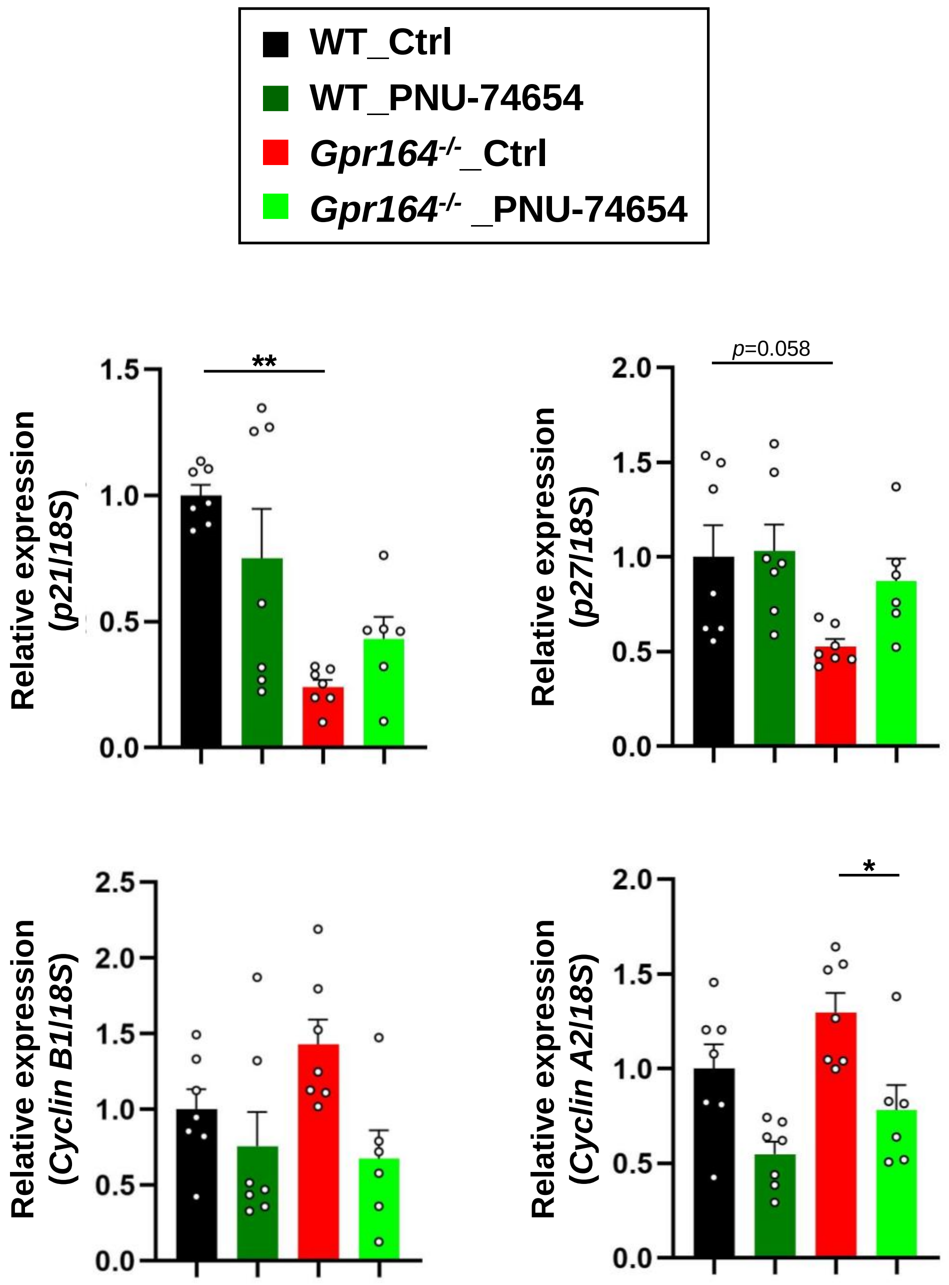


**Supplementary Figure 4**

| | gene | Forward | Reverse |
|---|---|---|---|
| human | *18S rRNA* | aaacggctaccacatccaag | cgctcccaagatccaactac |
| | *p21* | ggttcccgtttctccaccta | attgtgggaggagctgtgaa |
| | *p27* | ccggtggaccacgaagagt | gctcgcctcttccatgtctc |
| | *Cyclin D1* | ggactttgaggcaagtgtgg | tttcttcttgactggcacgc |
| | *Cyclin B1* | tgatactgcctctccaagcc | gctccatcttctgcatccac |
| | *Cyclin A2* | ggaccttcaccagacctacc | agtgtctctggtgggttgag |
| | *Occludin* | aagagttgacagtcccatggcatac | atccacaggcgaagttaatggaag |
| | *Claudin-3* | aaggtgtacgactcgctgct | gaagtcccggataatggtgtt |
| | *Zo-1* | tccgtgttgtggataccttg | ggatgatgcctcgttctacc |
| mouse | *18S rRNA* | acgctgagccagtcagtgta | acgctgagccagtcagtgta |
| | *Or51e1* | tgctgtgctaggtaacttgaca | ccagacaagcatcaaactggat |
| | *p21* | cccatacttccccttctgca | cccacttagtgtaccctgca |
| | *p27* | tcttcggcccggtcaa | ccggcagtgcttctccaa |
| | *Cyclin D1* | ctggccatgaactacctgga | atccgcctctggcattttgg |
| | *Cyclin B1* | agggtcgtgaagtgactggaaac | ttgggcacacaactgttctgc |
| | *Cyclin A2* | cacaacatgcccaacagtca | agtgtctctggtgggttgag |
| | *Occludin* | ggaccctgaccactatgaaacagatac | ataggtggatattccctgacccagtc |
| | *Zo-1* | cttctcttgctggccctaaac | tggcttcacttgaggtttctg |
| | *Lgr5* | ccacagcaacatcaggt | aacaaattggatggggttgt |
| | *Hes1* | gagtgcatgaacgaggtgac | cgttgatctgggtcatgcag |
| | *Atoh1* | gggtgagctggtaaggagaa | actacaaccccacccttcag |
| | *Tph1* | acatcagccgagaacagttg | aacgtcttccttcgcagtga |
| | *Muc2* | atgtcctgaccaagagcgaa | gacagtcttcaggcaggtct |
| | *Vil1* | cgtatcaagccgtcctgttg | ccctgataaaccaccatgcg |
| | *c-Myc* | agctggagatgatgaccgag | gaaggtctcgtcgtcaggat |
| | *Rtp1S* | tggtgcttattttgggccac | ccaccccaagcttagacaga |
| | *Rtp2* | gcctataccgatgcctccaa | cagaacaggcaccaacgaaa |
| | *Rtp3* | aggttccactgtccttcctg | ctcatcttcacccacccctt |
| | *Rtp4* | ggttccagtgttccagatgc | tgcgatttcaaagtgtccgg |

**Supplementary Table 1**
